# Electroencephalography for Radiation-Free Orientation Estimation of Directional Deep Brain Stimulation Leads

**DOI:** 10.1101/2024.02.27.582287

**Authors:** Suvadeep Maiti, Alan Bince Jacob, Marco Mancuso, Marie T. Krueger, Harith Akram, Kirill Aristovich, Vladimir Litvak

## Abstract

**Background:** Programming segmented deep brain stimulation (DBS) leads requires knowledge of their rotational orientation, but current postoperative workflows rely mainly on radiographic imaging.

**Objective:** To develop and evaluate a radiation-free electroencephalography (EEG) framework for directional lead orientation estimation, in a realistic phantom and postoperative human data.

**Methods:** A realistic head phantom containing a directional DBS lead was mounted on a rotating tray and stepped through known orientations in 5° increments to evaluate EEG-based orientation estimation. Stimulation artefact topographies were compared with forward-model predictions using Pearson correlation, and a tri-montage approach combined estimates from all three same-ring electrode pairs to improve accuracy. Robustness was tested by adding brain noise and reducing the electrode count. The method was then applied to a patient with bilateral directional DBS leads implanted in the globus pallidus. Robustness in the human data was assessed by reducing the sampling rate and the number of EEG electrodes.

**Results:** In the phantom, the best simplified forward model was a tangential BEM dipole positioned midway between the active contacts. The tri-montage reduced the maximal mean orientation error to 0.59° (95% CI: 4.48°). A conservative tray-position analysis yielded a pooled mean error of 2.05° (95% CI: 3.20°). In the human dataset, EEG-derived orientations differed from imaging-based references by about 8° for the right lead and 15° for the left lead, and remained stable after downsampling and montage reduction.

**Conclusions:** Phantom validation established the key modelling assumptions, and postoperative human recordings showed that EEG can recover clinically plausible directional lead orientations under realistic conditions.

## 1. Introduction

Directional DBS leads enable current steering and can widen the therapeutic window compared with conventional ring contacts by directing stimulation toward desired pathways and away from structures that mediate side effects [1, 2, 3]. In contrast to conventional cylindrical contacts, segmented leads introduce an additional rotational degree of freedom: the clinical effect of stimulation now depends not only on the contact level and stimulation parameters, but also on the azimuthal orientation of the active segment relative to surrounding anatomy. If the rotational orientation of the implanted lead is unknown, the anatomical direction faced by each segment remains uncertain, reducing the precision of image-guided programming.

This issue is clinically important because DBS targets are small, anatomically crowded structures in which modest changes in the direction of current spread can affect both therapeutic benefit and side-effect profile. Directional programming is intended to exploit this anatomical specificity, but its practical value depends on reliable knowledge of where each segment faces after implantation. Inaccurate orientation information can reduce confidence in postoperative programming decisions, complicate interpretation of stimulation-response relationships, and limit the usefulness of patient-specific modelling tools.

Current postoperative workflows determine lead orientation using CT, flat-panel CT, or rotational fluoroscopy and often require dedicated software to identify the radiographic marker artefact [4, 5, 6, 7, 8]. These methods are clinically useful and remain the practical reference standard in many settings. However, they expose patients to ionising radiation and depend on image quality and accurate interpretation of radiographic artefacts. They are also less suitable when repeated verification of lead orientation is required. This is relevant because directional leads can deviate from their intended implantation angle and may rotate after surgery [9, 10, 11]. A short, repeatable, radiation-free measurement would therefore have practical value, particularly during postoperative programming, longitudinal follow-up, or studies of lead stability over time.

Magnetoencephalography (MEG) has been proposed as a non-radiative alternative for localisation and orientation estimation of segmented DBS leads [12, 13]. However, MEG is expensive, not widely available in hospitals, and difficult to integrate into routine DBS care. EEG offers a more accessible option because it is inexpensive, common in neurology and neurophysiology settings, and compatible with short outpatient recordings. Each DBS pulse generates a brief stimulation artefact whose scalp topography depends on the position and orientation of the active contact pair, the conductive properties of the head, and the sensor montage. When stimulating between two segments on the same ring, the current direction is orthogonal to the lead resulting in scalp artefact topographies strongly dependent on the rotational angle. This was previously shown for MEG [13], but the same logic applies to EEG. Assuming that lead location can be measured with high precision from perioperative CT and MRI using established methods [14], lead orientation can be estimated by comparing the measured scalp topography with forward-model predictions across candidate angles.

This approach relies on two key assumptions. First, the stimulation artefact must contain sufficient orientation-specific information to discriminate between different segment directions. Second, a simplified forward model must capture this information with adequate fidelity to support estimation in empirical data.

In the present study, we tested these assumptions using a realistic head phantom, followed by validation in a human experiment.

The aims were threefold. First, to use the phantom to validate a simplified forward model and quantify estimator robustness under controlled conditions. Second, to test whether clinically plausible lead orientations can be recovered in a patient with bilateral pallidal directional leads. Third, to determine whether orientation estimates remain stable under conditions resembling a realistic device that could be used in an outpatient clinic, such as lower sampling rates and sparse sensor montages.

## 2. Methods

### 2.1. Phantom validation experiment

#### 2.1.1. Phantom setup and acquisition

Phantom validation was performed using a 3D-printed adult head tank developed for electrical impedance tomography experiments [15]. The phantom preserves realistic head geometry derived from CT and MRI, includes a perforated skull insert to regulate conductivity, and was filled with 0.9% isotonic saline to mimic the bulk conductive properties of the head volume conductor. Thirty-two EEG electrodes and one ground electrode were embedded in the outer wall according to the international 10–20 system [16]. A photograph of the setup is shown in Fig. 1A.

**Figure 1:**
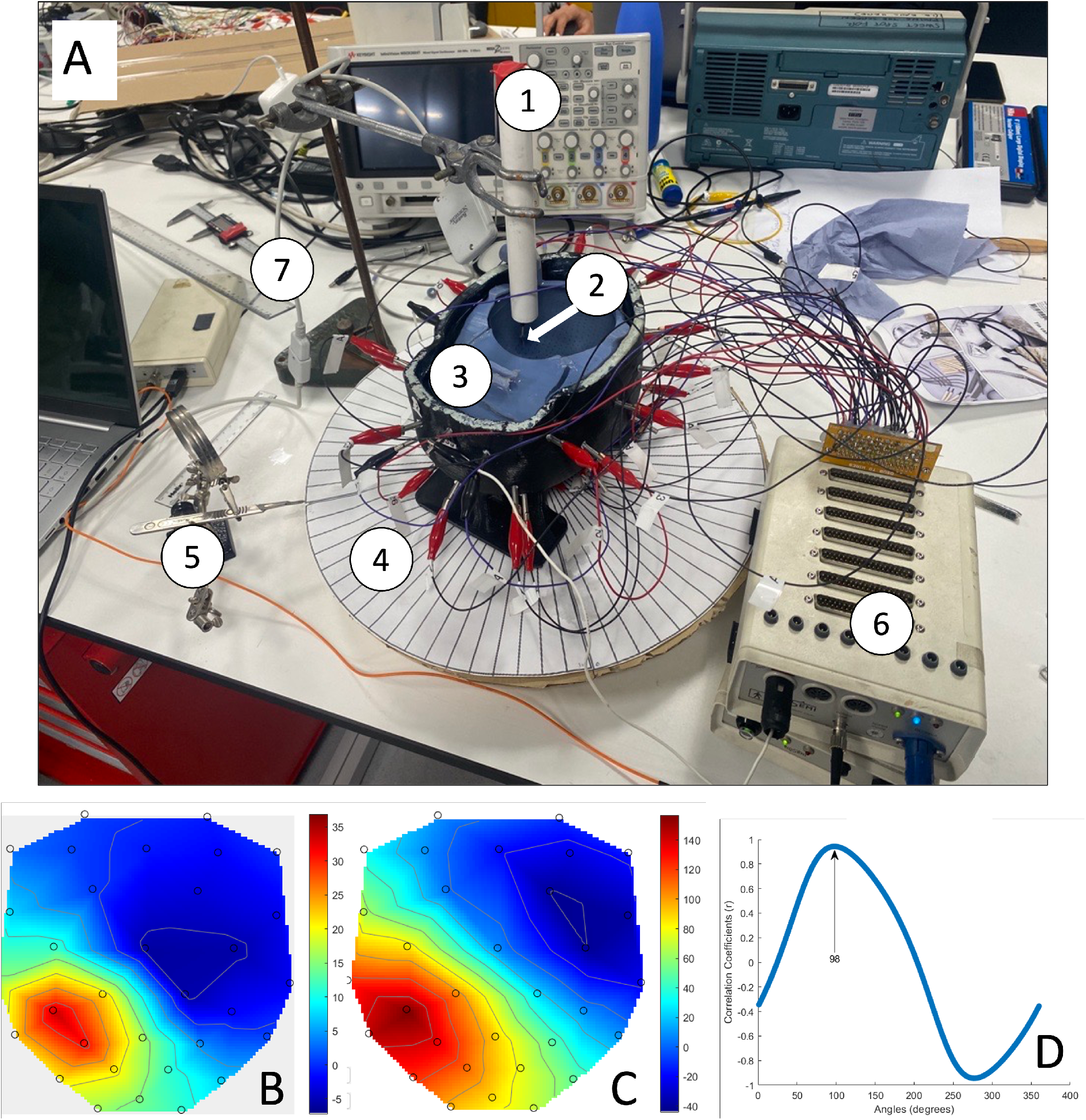
Phantom validation workflow. **(A)** Photo of the phantom setup. (1) electrode holder pipe; (2) electrode tip; (3) skull insert; (4) rotating tray; (5) current angle indicator; (6) EEG amplifier; (7) cable to current source. **(B)** Representative measured scalp topography from the phantom recording for one same-ring bipolar stimulation condition. **(C)** Corresponding simulated scalp topography from the forward model at the best-fitting orientation. **(D)** Pearson correlation as a function of rotation angle, with the peak indicating the estimated lead orientation.

A Boston Scientific Vercise Cartesia directional lead was positioned to approximate the location of DBS targets such as left subthalamic nucleus (STN) and globus pallidus internus (GPi), with the target coordinates scaled from the Colin27 template brain [17] and set manually with an electronic calliper. The phantom was fixed to a rotating tray marked at 5° intervals for controlled angular adjustments. Stimulation was delivered using a Keithley 6221 current source (Tektronix Inc., Beaverton, US) with clinically representative settings of 3 mA, 60 µs, and 130 Hz. EEG was acquired with a BioSemi system (Biosemi B.V., Amsterdam, the Netherlands) at 4096 Hz. The main analysis used the three same-ring bipolar stimulation pairs A^−^B^+^, B^−^C^+^, and C^−^A^+^. The head phantom was positioned on a rotatable platform with a fixed pointer used as the rotational reference. EEG recordings were acquired while stimulating the B^−^C^+^ same-ring bipolar pair. Starting from the reference position, the phantom was rotated in 5° increments over a 120° range, with one recording obtained at each orientation. Because the three segmented contacts are separated by 120°, these measurements were subsequently mapped to the equivalent orientations of the A^−^B^+^ and C^−^A^+^ pairs to represent the full 360° rotation.

#### 2.1.2. Phantom preprocessing and noise augmentation

Phantom EEG was analysed using the Statistical Parametric Mapping toolbox (SPM, https://www.fil.ion.ucl.ac.uk/spm/, [18]). An average-reference montage was applied across scalp electrodes. Signals were high-pass filtered above 120 Hz. The filtered recordings were segmented into epochs with a duration equal to the inverse of the stimulation frequency. Small timing drifts accumulated over the course of the recording, such that direct averaging of these epochs would have smeared the stimulation artefact peak and reduced spatial precision. To address this, a channel with a clear artefact was used to identify the stimulation-related peak within each epoch. A linear model of peak latency as a function of epoch number was then fitted and used to correct for latency shifts prior to averaging. This alignment ensured that stimulation artefacts from different trials were brought into a common temporal frame, sharpening the averaged response and improving the stability of the resulting scalp topographies. To estimate robustness in the presence of realistic noise, phantom averages were combined with human EEG from the publicly available mismatch negativity dataset distributed with SPM [19]. Random subsets of human epochs were averaged and added to the phantom artefact average across repeated iterations and increasing trial counts.

#### 2.1.3. FEM and BEM comparison

A detailed finite element (FEM) model was built in COMSOL (https://www.comsol.com/) using realistic lead geometry and phantom conductivity. Same-ring and diagonal segment configurations were explored to understand current flow and scalp field structure. In parallel, a boundary element (BEM) model was constructed using the Helsinki BEM framework [20]. The BEM source was defined as a dipole whose position and orientation were derived from the active segment pair. Comparing FEM and BEM predictions allowed us to determine whether the simplified BEM source retained the essential orientation information from the more detailed FEM representation.

#### 2.1.4. Phantom error metrics

Two error metrics were used. The first examined the difference between the tri-montage orientation estimate at pairs of lead orientations separated by 120°, exploiting the three-fold rotational symmetry of the segmented electrode; the expected difference is 120°. The second compared estimates at consecutive tray positions, differing by 5°. This metric provided a more conservative estimate because it also included mechanical imprecision of the rotating setup.

### 2.2. Human experiment

#### 2.2.1. Participant, recording, and stimulation protocol

The human dataset was acquired from a 54-year-old male patient with cervical dystonia implanted with bilateral directional DBS leads (Boston Scientific Vercise Cartesia) targeting the internal Globus Pallidus (GPi). EEG was recorded using a BrainVision actiCHamp Plus system (Brain Products GmbH, Gilching Germany) at 100 kHz. A passive cap was used to minimise delays between electrodes that could distort the artefact pattern. Scalp electrode impedances were kept below 10 kΩ. An auxiliary channel placed above the implanted pulse generator captured a combined ECG and stimulation reference signal. Stimulation was delivered continuously at 130 Hz with biphasic pulses of 1 mA, 2 mA, 3 mA, and 4 mA. Pulse width was 60 µs for the left GPi and 90 µs for the right GPi. For each hemisphere, the three same-ring bipolar segment-pair configurations A^−^B^+^, B^−^C^+^, and C^−^A^+^ were tested. Each condition was recorded for approximately 1 min.

#### 2.2.2. Electrode localisation and imaging-informed anatomy

To obtain subject-specific sensor positions, the participant’s head surface and EEG cap were scanned using a handheld 3D scanner (EinScan H1, SHINING 3D Technology GmbH, Stuttgart, Germany) after electrode placement. Electrode coordinates were co-registered to the structural MRI using fiducial landmarks. Postoperative CT and preoperative MRI were used to localise the lead in Lead-DBS (https://www.lead-dbs.org/) [14] using the default pipeline. This provided the coordinates of the segmented contacts and the direction of the lead’s longitudinal axis (i.e., the axis running along the length of the DBS lead), which were required for forward modelling.

#### 2.2.3. Human data preprocessing and topography extraction

Data were processed using SPM and custom MATLAB scripts similarly to the phantom data described above.

After drift correction, epochs belonging to the same stimulation condition were averaged to obtain a condition-specific artefact waveform (Fig. 2C). The averaged response contained a sharp, time-locked stimulation artefact, followed by a second closely spaced deflection. The measured scalp topography was extracted from a fixed 0.2 ms window, positioned on the initial artefact complex. This topography was compared with forward-model predictions across candidate ring-plane orientations, and the orientation was estimated as the angle of maximum absolute correlation.

**Figure 2:**
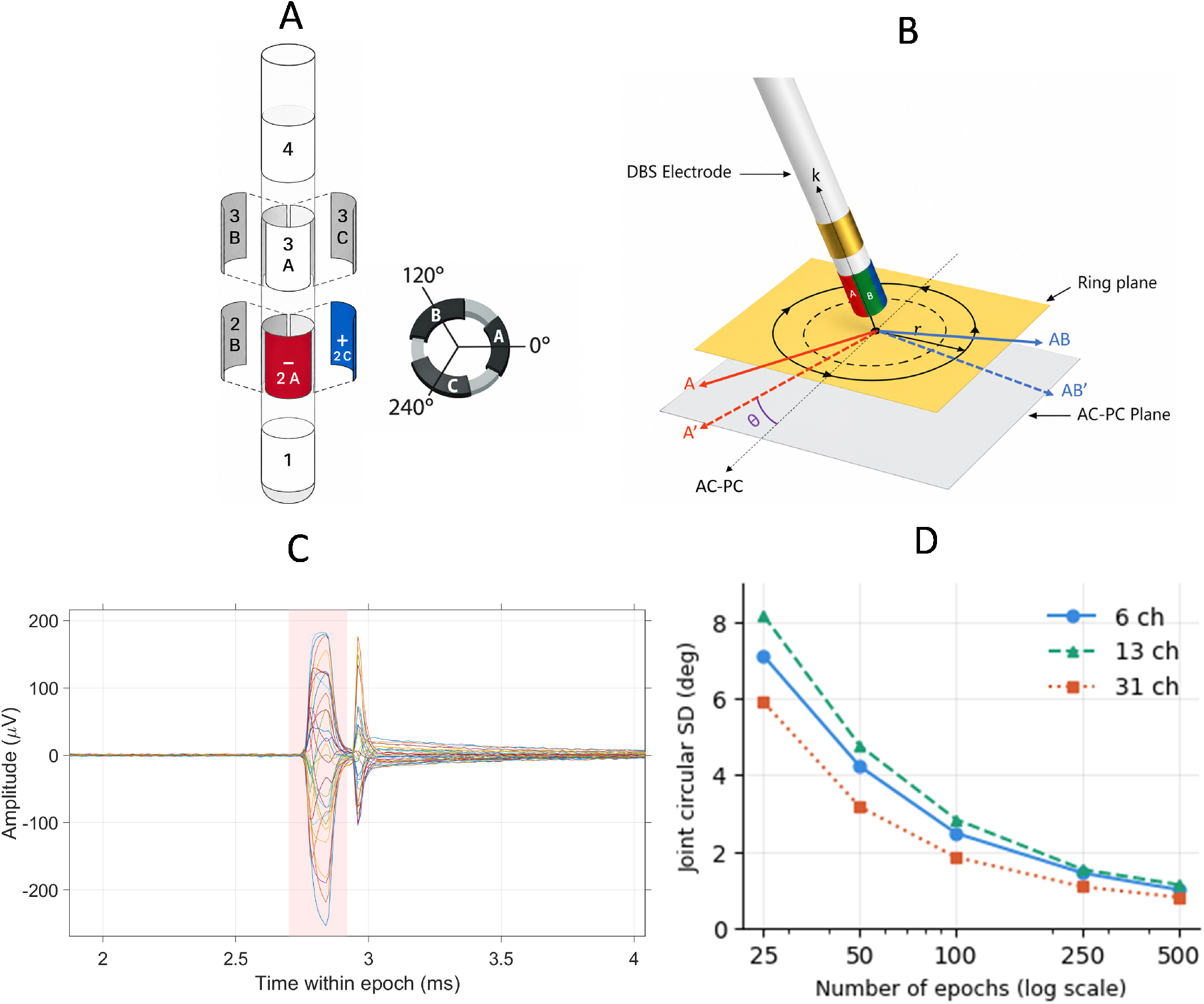
Human DBS orientation estimation overview. **(A)** Directional DBS lead with three segmented contacts (A, B, C) and an example same-ring bipolar montage (A^−^B^+^). **(B)** Geometry and angle convention: *k* denotes the lead-shaft axis; the ring plane is orthogonal to *k*. The AC–PC plane defines the anatomical reference; the segment-A direction (shown as *A*) is projected onto the AC–PC plane (*A*^′^), and θ is the in-plane orientation relative to AC–PC. **(C)** Stimulation artefact averaged across 100 epochs, shown for all EEG channels (one trace per channel). The shaded band marks the time window whose mean scalp topography was used for orientation estimation. **(D)** Robustness summary showing absolute orientation error under montage reduction (e.g., outer-ring electrodes), illustrating that the estimate is primarily driven by stable spatial topography.

#### 2.2.4. Subject-specific BEM forward model

A three-layer BEM head model [21] was generated from the participant’s MRI, comprising scalp, skull, and brain compartments. Co-registered electrode positions were combined with the BEM surfaces. For each active contact pair, an equivalent current dipole was placed at the midpoint between the contacts. The dipole orientation was rotated through 360 candidate angles within the ring plane perpendicular to the DBS lead shaft (Fig. 2 A,B) using Eq. 1.

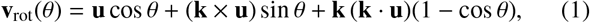

where **k** is the unit vector along the lead axis, **u** is the reference direction in the ring plane, and θ is the candidate orientation angle.

The projection of the anterior commissure–posterior commissure direction onto the rotation plane defined the anatomical zero-angle reference (Fig. 2B).

#### 2.2.5. Orientation estimation from measured and simulated scalp topographies

For each bipolar stimulation montage *m* ∈ {*AB, BC, CA*}, the DBS artefact was extracted from the average across a series of stimuli. For each candidate angle θ ∈ [0°, 360°), a simulated scalp topography was computed using the contact-pair-specific BEM lead-field matrix,

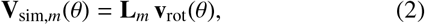

where 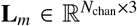 is the BEM lead-field for montage *m* and 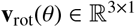 is the dipole orientation vector rotated within the ring plane.

Similarity between the measured and simulated topographies was quantified with Pearson correlation,

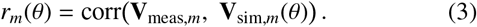

The per-montage orientation estimate was obtained as the angle maximising the absolute correlation,

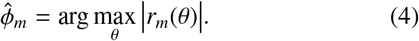

The 180° ambiguity was then resolved by the sign of the correlation at the peak: if 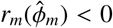, the estimate was shifted by

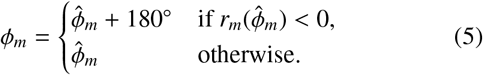

#### 2.2.6. Tri-montage combination

The three same-ring bipolar montages *AB, BC*, and *CA* subtend known geometric offsets δ_*m*_ relative to a common reference direction. Each per-montage estimate was therefore corrected for its offset,

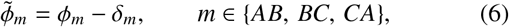

and the three offset-corrected estimates were combined using the axial circular mean,

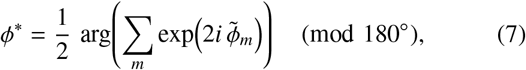

which correctly accounts for the 180° periodicity of an axial quantity. The final lead orientation ϕ^∗^ was then projected into the anatomical AC–PC reference frame for reporting.

#### 2.2.7. Robustness to sampling rate and electrode reduction

To assess feasibility and reproducibility under acquisition conditions more representative of a simplified clinical system, the tri-montage orientation estimate was re-evaluated across a grid of sampling rates, electrode configurations, and epoch counts.

The native EEG recordings were acquired at 100 kHz. To simulate recordings obtained using lower-sampling-rate acquisition hardware, an anti-alias low-pass filter was applied before resampling to 50 kHz, 10 kHz, and 5 kHz. This step attenuated frequency components above the usable bandwidth of each target sampling rate before decimation, preventing frequencies above the target Nyquist frequency from folding into the retained spectrum. The resulting signals therefore represented bandwidth-limited lower-rate recordings rather than direct unfiltered decimation of the 100 kHz data. Anti-alias filtering used an eighth-order Butterworth low-pass filter with cutoffs of 20, 4, and 2 kHz before downsampling to 50, 10, and 5 kHz, respectively. The electrode montage was independently reduced from the full 31-channel configuration to 13- and 6-channel subsets (as in Fig. 3A). The 13-channel subset comprised Fp1, Fp2, F7, F8, T7, T8, TP9, TP10, P7, P8, O1, O2, and Iz; the 6-channel subset comprised Fp1, Fp2, T7, T8, O1, and O2.

**Figure 3:**
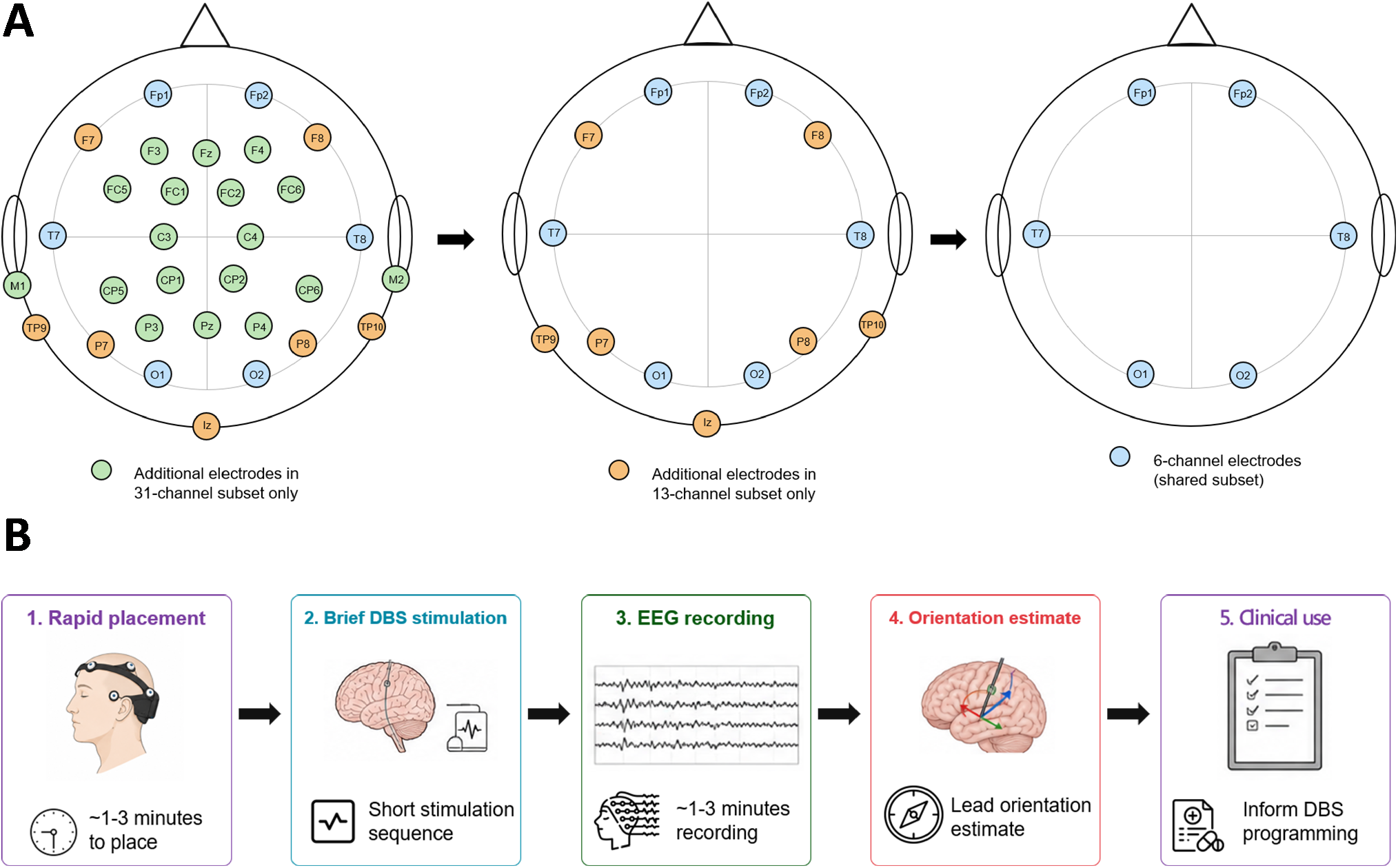
Electrode configurations and proposed clinical workflow for EEG-based DBS lead orientation estimation. **(A)** Electrode configurations evaluated in the human robustness analysis. The full 31-channel montage was progressively reduced to 13- and 6-channel subsets. Blue electrodes indicate the 6-channel shared subset, orange electrodes indicate the additional electrodes retained in the 13-channel subset, and green electrodes indicate electrodes present only in the full 31-channel configuration. **(B)** Proposed clinical workflow comprising rapid electrode placement, brief same-ring bipolar DBS stimulation, short EEG recording, orientation estimation using subject-specific forward modelling, and use of the resulting orientation estimate to support directional DBS programming.

For each combination of sampling rate and electrode subset, the estimate was repeated for epoch counts of 25, 50, 100, 250, and 500. At every grid point, the orientation was re-estimated on 500 random epoch subsets, and reproducibility was quantified as the circular (axial, modulo 180°) standard deviation of the tri montage-wise orientation estimate across these subsets, together with its 95th-percentile-trimmed value to discount occasional outliers.

## 3. Results

### 3.1. Phantom validation

#### 3.1.1. A tangential BEM dipole captures the key orientation information from the FEM model

The phantom experiments first tested whether a simplified BEM representation could preserve the orientation-dependent information observed in the more detailed FEM simulations. In the representative example shown in Fig. 1B–D, same-ring bipolar stimulation produced distinct scalp potential patterns that varied systematically with lead orientation. FEM simulations indicated that these same-ring bipolar pairs generated predominantly circumferential current flow, whereas diagonal configurations introduced a stronger vertical current component and were less clearly separable. When FEM-derived scalp maps were compared with BEM predictions, the closest agreement was obtained when the BEM source was modeled as a tangential dipole positioned midway between the active contacts. Under this simplified representation, the BEM reproduced representative FEM orientations such as 60°, 120°, and 180°, supporting its use as the main forward model for orientation estimation.

#### 3.1.2. The tri-montage approach markedly improves phantom performance

After selecting the tangential BEM dipole as the simplified forward model, the phantom experiment was used to quantify orientation-estimation performance under controlled conditions. Exploiting the three-fold rotational symmetry of the same-ring configuration, the phantom was stepped through orientations in 5° increments to provide estimates at 72 unique orientations spanning the full 360°.

Using a single same-ring bipolar pair with approximately 1 000 averaged trials yielded a maximal absolute mean error of 14.99°, with a pooled mean error of 6.10° across all tested orientations (Fig. 4A). The magnitude of this error varied systematically with lead orientation, indicating that geometry-dependent bias dominated over random noise.

**Figure 4:**
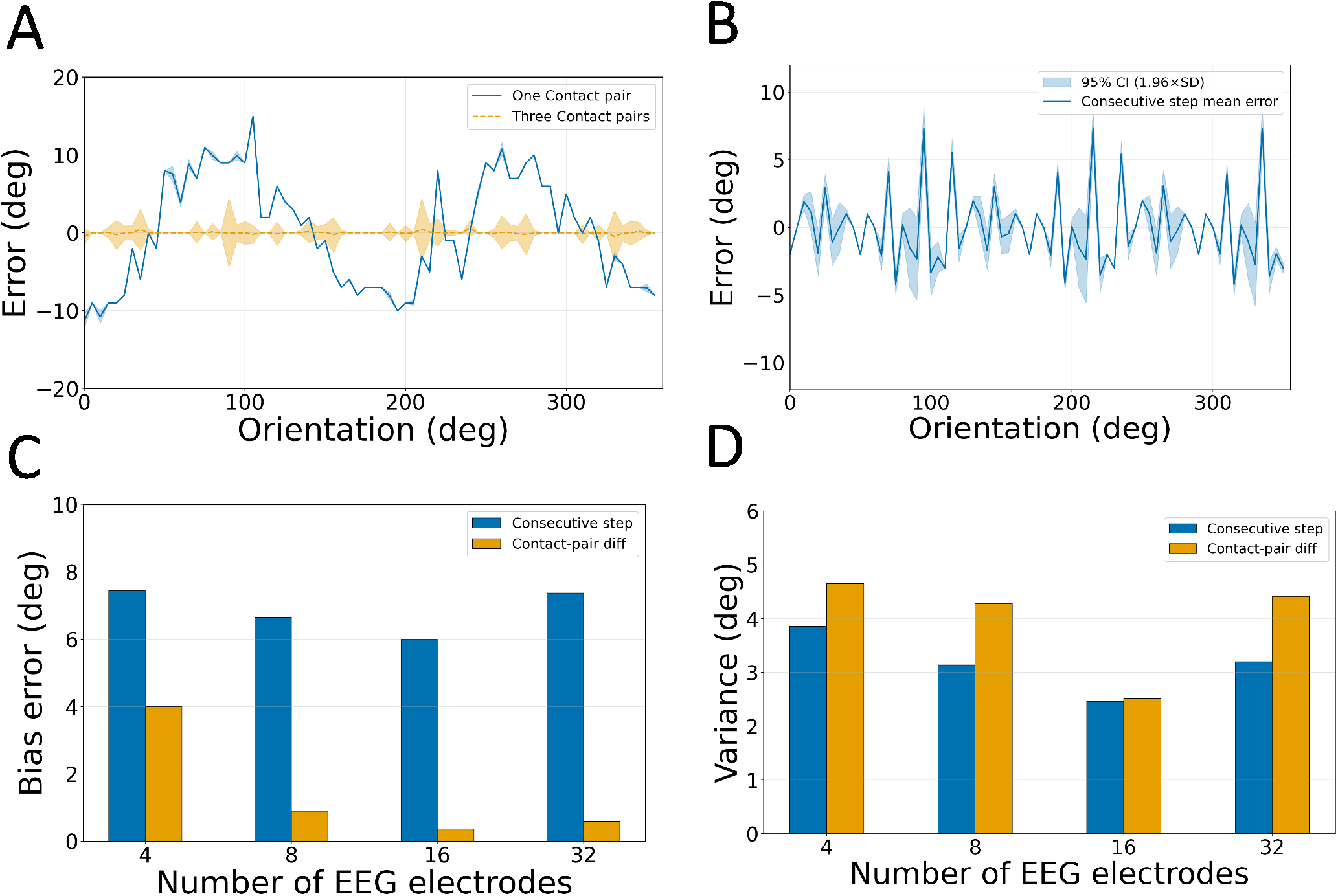
Phantom performance summary using the tri-montage method. **(A)** Measurement errors for contact-pair differences (expected difference: 120°), shown as a function of lead orientation when using one contact pair (blue line) and three pairs combined via tri-montage (orange line). The shaded area represents the 95% confidence interval (1.96 ×SD) derived from 32 repetitions with different trial subsets and noise realisations. **(B)** Measurement errors for the difference between consecutive turntable positions (expected difference: 5°), shown as a function of lead orientation using the three pairs combined. The shaded area indicates the 95% confidence interval (1.96 ×SD) from 32 repetitions with different trial subsets and noise realisations. **(C)** Errors due to bias. Bars show the maximal mean error across orientations for estimating orientation differences between consecutive turntable positions (blue bars; cf. panel B) and between two contact pairs using tri montage (orange bars; cf. panel A, orange line), for different numbers of EEG electrodes. **(D)** Errors due to measurement variance. Bars show the maximal 95% confidence interval (1.96×SD over 32 repetitions). Bar colors and x-axis categories correspond to panel C.

Combining all three same-ring bipolar pairs produced a marked improvement (Fig. 4A). This tri-montage approach reduced the maximal mean error to 0.59°, with a maximum 95% confidence interval of 4.48° across all 72 tested orientations.

#### 3.1.3. Robustness to reduced channel count supports clinic-friendly acquisition

The phantom data were also used to assess robustness to reduced channel counts. As shown in Fig. 4C,D, estimation errors increased for sparser electrode configurations, but the tri-montage method remained substantially more accurate than the single-pair approach at every channel count tested. With 16 channels the tri-montage maximal mean contact-pair error was 0.38° (95% CI: 2.52°), and with 8 channels it was 0.88° (95% CI: 4.27°). Even the most reduced four-electrode configuration retained a tri-montage maximal mean contact-pair error of 4.00° (95% CI: 4.65°), suggesting the method does not strictly require a dense electrode array. The consecutive-step error (blue bars, Fig. 4C,D) followed a similar trend: maximal mean errors were 6.00°, 6.66°, and 7.44° for 16, 8, and 4 electrodes respectively, with corresponding 95% confidence intervals of 2.46°, 3.14°, and 3.86°.

Acceptable performance—defined here as a maximal mean contact-pair error below 1°—was retained with as few as 8 electrodes distributed around the head. Although precision degraded relative to the full 32-channel configuration, this finding indicates that the tri-montage method does not depend on dense-array EEG and could remain feasible in simplified, clinic-friendly acquisition settings.

### 3.2. Human experiment

#### 3.2.1. Measured and simulated topographies preserve the same-ring geometry

Having established the modelling assumptions in the phantom, we then applied the same framework to the human recordings. The patient data showed clear, reproducible stimulation artefact topographies for all three same-ring segment-pair configurations in both hemispheres. When each measured scalp map was compared with the corresponding BEM predictions across 360 candidate angles, distinct correlation maxima were observed rather than broad, poorly constrained plateaus (Fig. 4C). This indicated that the measured EEG contained reliable orientation information even in the presence of real postoperative anatomy, physiological noise, and hardware artefacts.

**Table 1:** Key phantom-validation results.

| Metric | Value |
| --- | --- |
| Simplified forward model | Tangential BEM dipole |
| Single-pair max. mean error | 14.99° |
| Single-pair mean error (all orientations) | 6.10° |
| Tri-montage max. mean error | 0.59° |
| Tri-montage 95% confidence interval | 4.48° |
| Smallest montage (max. error <1°) | 8 electrodes |

**Table 2:** Human postoperative orientation results.

| Side | EEG estimate<br>(° left of AC-PC) | CT/MRI estimate<br>(° left of AC-PC) | Difference |
| --- | --- | --- | --- |
| Right GPi | 50.1 | 58 | 8° |
| Left GPi | 19.6 | 35 | 15° |

For the right GPi lead, the peak ring-plane angles were 210°, 311°, and 79° for A^−^B^+^, B^−^C^+^, and C^−^A^+^, with peak correlations of 0.98, 0.91, and 0.98. For the left GPi lead, the peak angles were 157°, 286°, and 56°, with peak correlations of 0.95, 0.92, and 0.88 (as shown in Fig. 5C & Supplementary Tab. T1).

**Figure 5:**
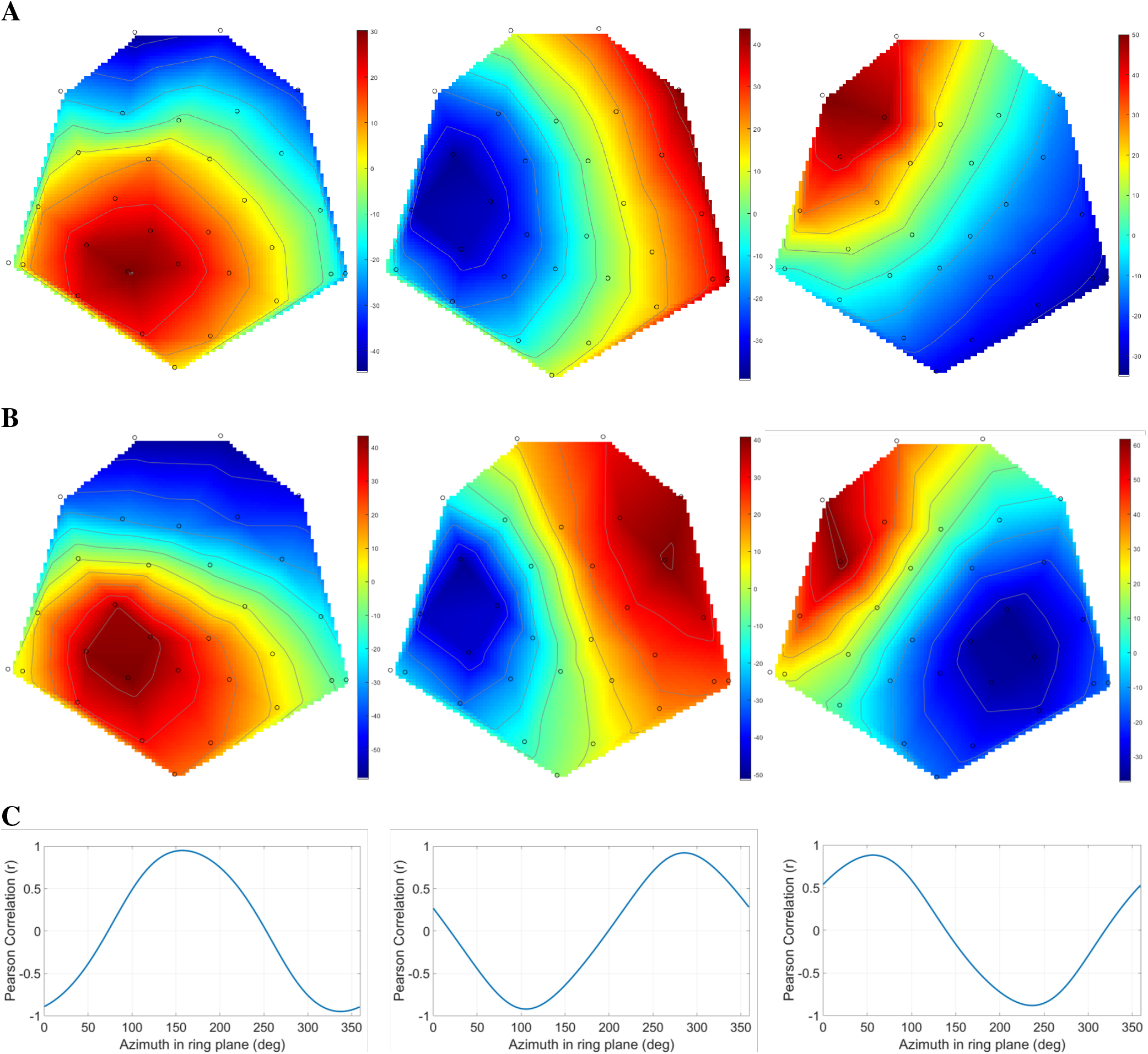
Patient results and orientation predictions for the left GPi. **(A)** Measured scalp topographies for A^−^B^+^, B^−^C^+^, and C^−^A^+^, shown from left to right. **(B)** Simulated scalp topographies at the corresponding best-fitting ring-plane angles. **(C)** Pearson correlation (*r*) as a function of ring-plane angle for the three same-ring bipolar stimulation pairs. Peak correlations occurred at 157° for A^−^B^+^ (*r* = 0.95), 286° for B^−^C^+^ (*r* = 0.92), and 56° for C^−^A^+^ (*r* = 0.88). Together, the panels demonstrate close agreement between the measured EEG topographies and the BEM predictions and recover the expected same-ring geometry.

In both hemispheres, the three best-fit solutions approximately preserved the expected 120° spacing between same-ring segment pairs (Supplementary Tab. T2), supporting the internal consistency of the patient data.

#### 3.2.2. EEG-derived lead orientations agree with imaging references

The orientation estimates obtained from the three bipolar montages were adjusted using the known 1 20° spacing between segments so that each represented the orientation of segment A. The adjusted estimates were combined using a circular mean and then expressed relative to the AC–PC line. For the right lead, the resulting EEG-based orientation was 50.1° left of the AC–PC line. The corresponding CT/MRI reference indicated about 58° left, so the disagreement was approximately 8°. For the left lead, the EEG estimate was 19.6° left of AC–PC, compared with an imaging-based reference of about 35° left, corresponding to a difference of about 15°.

#### 3.2.3. Robustness to anti-aliased sampling-rate and electrode reduction

A major translational question was whether the method depended on ultra-high-rate acquisition and a full EEG cap. The insights from the patient data were encouraging. Across the robustness grid, the tri-montage orientation estimate was highly reproducible under random epoch subsampling, and its reproducibility improved steadily with the number of available epochs: the median circular standard deviation fell from 5.5° at 25 epochs to 0.8° at 500 epochs (Supplementary Table T3). At the fullest sampling (500 epochs) the orientation was recovered with a circular SD below ∼ 1.1° across every condition, and the 95th-percentile-trimmed values were essentially unchanged, indicating that this spread reflects genuine sampling variability rather than occasional outliers.

This stability was largely preserved as acquisition was made more realistic (Fig. 2D). Following anti-aliased resampling of the native 100 kHz recordings, reducing the sampling rate to 50 kHz, 10 kHz, and 5 kHz produced only modest changes in the reproducibility of the orientation estimate. Similarly, restricting the montage from the full 31-channel configuration to 13- and 6-channel subsets only modestly increased variability (median circular SD 1.7° for the full set versus approximately 2.2° for the reduced subsets at 500 epochs).

To verify that anti-aliased resampling preserved the spatial structure of the stimulation artefact, we additionally examined a representative recording using an average of 100 epochs. Relative to the native 100 kHz topography, spatial correlation remained high after resampling to 50 kHz (*r* = 0.99) and 10 kHz (*r* = 0.99), and decreased to *r* = 0.97 at 5 kHz. The corresponding RMSE values were 1.84, 11.56, and 24.96 µV, respectively. The relative amplitude scale was 0.964, 0.774, and 0.505 at 50 kHz, 10 kHz, and 5 kHz, respectively. This representative analysis illustrates that anti-aliased resampling largely preserved the spatial topography at 50 kHz and 10 kHz, while greater attenuation was observed at 5 kHz.

The angular geometry underlying the tri-montage fit also remained coherent under these reduced conditions. As an additional check on the stability of the orientation-relevant artefact pattern, we compared averaged responses across stimulation amplitudes. Increasing the stimulation current primarily increased the artefact magnitude, while the temporal waveform and relative channel-wise distribution remained similar (Supplementary Fig. S1). This behaviour is consistent with the correlation-based orientation estimator, which relies on the relative spatial topography rather than the absolute signal amplitude.

## 4. Discussion

This manuscript is organised around the progression from controlled validation to translational testing. The phantom experiments established three important points before any patient inference was attempted: first, that same-ring bipolar segment pairs produce the clearest orientation-sensitive scalp patterns; second, that a tangential midpoint dipole is a defensible simplification of the full FEM source; and third, that pooling across the three same-ring pairs improves estimator stability. Those findings justify the simplified BEM framework that was then applied to the human data.

For context, previous phantom studies of directional DBS lead orientation have reported errors on the order of a few degrees using established imaging-based approaches. The CT-based DiODe algorithm achieved an orientation accuracy of− 0.6°± 1.5° relative to ground truth [8], while Sitz et al. reported a mean deviation of −0.3°± 1.9° for CT when the lead polar angle was ≤40°, with substantially larger errors for more oblique trajectories [6]. Radiation-free MEG approaches have reported orientation errors of approximately 11° in the initial SQUID-MEG implementation and approximately 4° with the subsequent MaDoPO approach [13, 12]. Although differences in phantom geometry, acquisition protocols, and error definitions preclude direct numerical comparison, the present EEG phantom results are of a comparable order of magnitude to previously reported imaging- and MEG-based orientation-estimation methods under controlled conditions.

Against that validated background, the postoperative human experiment becomes easier to interpret. Subject-specific EEG recovered orientation-dependent scalp topographies from both GPi leads, preserved the expected same-ring segment geometry, and produced final lead orientations that were reasonably close to imaging-based references. The right-sided solution differed from CT/MRI by about 8° and the left-sided solution by about 15°. For a single-patient feasibility dataset that includes real anatomy, physiological noise, postoperative lead localisation uncertainty, and simplified source modelling, that level of agreement is encouraging.

The human orientation estimate remained stable when the data were downsampled from the native 100 kHz to 10 and 5 kHz and when the montage was reduced from the full electrode set to sparser 13- and 6-channel subsets. These findings indicate that the critical information is carried by the spatial pattern of the stimulation artefact rather than by the full temporal precision of the original acquisition. In practical terms, this implies that the method could potentially be implemented on more widely available clinical EEG systems, and with a far simpler setup, than the research-grade recording used here, since a future clinic-friendly implementation may not require ultra-high-rate acquisition or a full research EEG cap.

Taken together, the phantom and human results suggest a coherent workflow: routine postoperative imaging provides the lead location, while a short EEG recording supplies an artefact topography that is matched against subject-specific forward-model predictions to infer the lead’s rotational orientation. In this sense, EEG does not replace imaging but complements it with a radiation-free readout that can be repeated during outpatient programming or longitudinal follow-up whenever uncertainty about lead orientation remains.

### 4.1. Limitations

Several limitations should be considered. First, the phantom setup was subject to mechanical positioning uncertainty due to the rotation tray, which may have increased the measured orientation error and therefore resulted in a conservative estimate of the achievable accuracy. The method also relies on accurate postoperative lead localisation and EEG sensor coordinates. These requirements are compatible with current clinical and experimental workflows, where postoperative CT or MRI is routinely used to verify lead placement and optical methods can provide accurate EEG electrode localisation [22, 23, 24].

Second, the imaging-based lead orientation used as the reference cannot be regarded as a perfect ground truth because CT-based orientation estimates have finite precision. In addition, the CT used as the reference was acquired at the time of lead implantation, whereas the EEG recording was performed over a year later. Consequently, delayed lead rotation between the postoperative CT and EEG recording cannot be excluded. Future validation could use photon-counting CT [25] as a more accurate imaging-based reference, owing to its improved spatial resolution and reduced metal artefacts compared with conventional CT.

Differences in STN lead trajectory, including variation in both the horizontal and vertical implantation angles, may influence the orientation estimation error. Validation in a larger cohort with a wider range of lead trajectories will therefore be an important future step.

Finally, the present approach requires stimulation between segmented contacts on the same ring. It is therefore applicable only to DBS systems that support same-ring bipolar stimulation, which would exclude, for instance, the current Medtronic stimulators.

## 5. Conclusions and Future Work

A subject-specific EEG framework can recover clinically plausible orientations of directional DBS leads in postoperative human recordings, while maintaining stable orientation estimates under anti-aliased reductions in sampling rate and sparse electrode montages. Phantom validation supports the simplified forward model, the use of same-ring bipolar stimulation, and the robustness gained by combining estimates from the three segment pairs. Together, these findings support EEG as a promising radiation-free adjunct for postoperative directional lead orientation verification.

A future system could use a small number of strategically positioned scalp electrodes, brief DBS stimulation, and a short EEG recording followed by automated orientation estimation (Fig. 3B). The reduced 13- and 6-channel configurations evaluated here (Fig. 3A), together with the sampling-rate analyses, suggest that a compact and rapidly deployable system may be feasible without requiring the full research-grade EEG setup used in this study. Future work should prospectively evaluate dedicated reduced-electrode and lower-sampling-rate hardware in a larger patient cohort and determine the optimal balance between electrode number, sampling rate, setup time, and orientation accuracy for routine clinical use.

## Ethics statement

The patient study was approved by the Bradford Leeds Research Ethics Committee, and the participant provided written informed consent prior to study participation.

## Acknowledgements

We thank the teams who developed the UCL head phantom and SPM analysis tools, as well as the clinical and technical teams whose infrastructure supported both the phantom and human recordings. The Functional Neurosurgery Unit is supported by the National Institute for Health and Care Research University College London Hospitals Biomedical Research Centre.

## Data availability

The code and phantom data used in this study are publicly available at https://github.com/suvadeepmaiti/seg_lead. Human participant data are available from the corresponding author upon reasonable request, subject to applicable ethical and privacy restrictions.

## CRediT authorship contribution statement

Suvadeep Maiti: Conceptualization, Methodology, Investigation, Software, Formal analysis, Visualization, Writing – original draft. Alan Bince Jacob: Methodology, Investigation, Software, Formal analysis, Visualization, Writing. Marco Mancuso: Investigation, Writing – review & editing. Marie T. Krueger: Methodology, Investigation, Writing – review & editing. Harith Akram: Clinical input, Methodology, Writing – review & editing. Kirill Aristovich: Supervision, Methodology, Writing – review & editing. Vladimir Litvak: Conceptualization, Supervision, Investigation, Formal analysis, Writing.

