## Supplementary material for "Electroencephalography for Radiation-Free Orientation Estimation of Directional Deep Brain Stimulation Leads"

### Supplementary Tables

Table T1: Peak angles and peak correlations for same-ring bipolar configurations in the human GPi leads.

| **Side** | **Bipolar pair** | **Peak angle** | **Peak correlation,** $r$ |
|---|---|---|---|
| Right GPi | A–B+ | 210° | 0.978 |
| Right GPi | B–C+ | 311° | 0.910 |
| Right GPi | C–A+ | 79° | 0.978 |
| Left GPi | A–B+ | 157° | 0.949 |
| Left GPi | B–C+ | 286° | 0.921 |
| Left GPi | C–A+ | 56° | 0.881 |

Table T2: Pairwise angular separations between same-ring bipolar configurations, computed from the ring-plane best angles.

| **Side** | **Comparison** | **Circular separation rule** | $\Delta\theta$ |
|---|---|---|---|
| Right GPi | AB vs BC | $\lvert 311 - 210 \rvert$ | 101° |
| Right GPi | BC vs CA | $360 - \lvert 311 - 79 \rvert$ | 128° |
| Right GPi | CA vs AB | $\lvert 210 - 79 \rvert$ | 131° |
| Left GPi | AB vs BC | $\lvert 286 - 157 \rvert$ | 129° |
| Left GPi | BC vs CA | $360 - \lvert 286 - 56 \rvert$ | 130° |
| Left GPi | CA vs AB | $\lvert 157 - 56 \rvert$ | 101° |

*Note.* Circular separations were computed on a 0°–360° ring. The expected same-ring spacing is approximately 120°; deviations reflect measurement noise, forward-model uncertainty, and subject-specific recording conditions.

Table T3: Reproducibility of the tri-montage orientation estimate under random epoch subsampling. The mean is essentially invariant across conditions, while the SD decreases with more epochs and increases modestly with fewer electrodes or lower sampling rate.

| **Electrodes** | **Sampling rate** | **Number of epochs per subset** | | | | |
|---|---|---|---|---|---|---|
| | | **25** | **50** | **100** | **250** | **500** |
| 31 ch | 100 kHz | $49.2 \pm 5.91$ | $49.6 \pm 3.17$ | $49.2 \pm 1.85$ | $49.2 \pm 1.07$ | $49.2 \pm 0.80$ |
| 31 ch | 10 kHz | $49.4 \pm 4.75$ | $49.2 \pm 2.63$ | $49.2 \pm 1.61$ | $49.2 \pm 0.95$ | $49.1 \pm 0.70$ |
| 31 ch | 5 kHz | $49.3 \pm 3.46$ | $49.2 \pm 2.12$ | $49.2 \pm 1.39$ | $49.2 \pm 0.84$ | $49.3 \pm 0.64$ |
| 13 ch | 100 kHz | $50.4 \pm 8.18$ | $50.1 \pm 4.77$ | $49.9 \pm 2.84$ | $49.6 \pm 1.52$ | $49.7 \pm 1.13$ |
| 13 ch | 10 kHz | $50.6 \pm 6.05$ | $49.7 \pm 3.33$ | $49.6 \pm 2.06$ | $49.8 \pm 1.36$ | $49.7 \pm 0.90$ |
| 13 ch | 5 kHz | $50.1 \pm 4.89$ | $49.9 \pm 2.67$ | $50.0 \pm 1.90$ | $50.0 \pm 1.13$ | $49.9 \pm 0.78$ |
| 6 ch | 100 kHz | $46.3 \pm 7.12$ | $46.6 \pm 4.23$ | $46.6 \pm 2.48$ | $46.4 \pm 1.43$ | $46.4 \pm 0.99$ |
| 6 ch | 10 kHz | $46.8 \pm 5.44$ | $46.7 \pm 3.26$ | $46.6 \pm 2.13$ | $46.6 \pm 1.19$ | $46.6 \pm 0.92$ |
| 6 ch | 5 kHz | $46.9 \pm 3.87$ | $46.8 \pm 2.60$ | $46.9 \pm 1.65$ | $46.9 \pm 1.08$ | $46.9 \pm 0.71$ |

*Note. Each cell gives the mean ± circular (axial) standard deviation over 500 random epoch subsets, for the right GPi lead.*

### Supplementary Figures

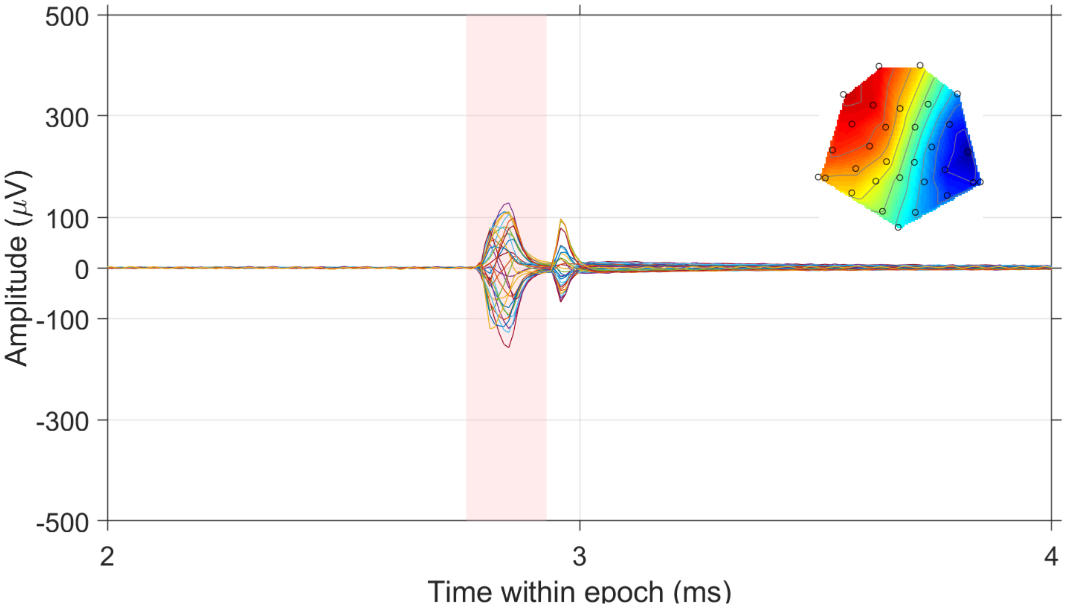


(a) 3 mA

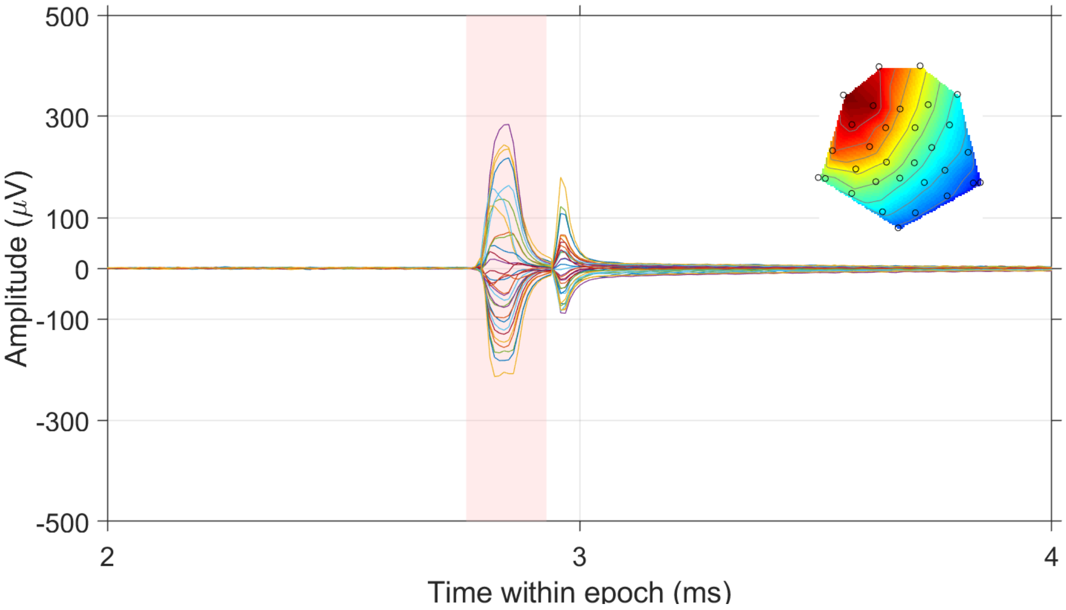


(b) 4 mA

Figure S1: Stimulation artefact at two stimulation amplitudes. Artefact averaged across 100 epochs and shown for all EEG channels (one trace per channel) at (a) 3 mA and (b) 4 mA. The artefact amplitude scales with stimulation current while its temporal shape and channel-wise spatial pattern are preserved, indicating that the orientation-relevant topography is stable across the tested amplitudes.
